# Antigen-specific humoral immune responses by CRISPR/Cas9-edited B cells

**DOI:** 10.1101/530832

**Authors:** Harald Hartweger, Andrew T. McGuire, Marcel Horning, Justin J. Taylor, Pia Dosenovic, Daniel Yost, Anna Gazumyan, Michael S. Seaman, Leonidas Stamatatos, Mila Jankovic, Michel C. Nussenzweig

## Abstract

A small number of HIV-1 infected individuals develop broadly neutralizing-antibodies to the virus (bNAbs). These antibodies are protective against infection in animal models. However, they only emerge 1 - 3 years after infection, and show a number of highly unusual features including exceedingly high levels of somatic mutations. It is therefore not surprising that elicitation of protective immunity to HIV-1 has not yet been possible. Here we show that mature, primary mouse and human B cells can be edited *in vitro* using CRISPR/Cas9 to express mature bNAbs from the endogenous *Igh* locus. Moreover, edited B cells retain the ability to participate in humoral immune responses. Immunization with cognate antigen in wild type mouse recipients of edited B cells elicits bNAb titers that neutralize HIV-1 at levels associated with protection against infection. This approach enables humoral immune responses that may be difficult to elicit by traditional immunization.

**One-sentence summary:** B cells edited by CRISPR/Cas9 to produce antibodies participate in humoral immune reactions and secrete neutralizing serum titers of anti-HIV bNAbs.

---

Although a vaccine for HIV remains elusive, anti-HIV-1 bNAbs have been identified and their protective activity has been demonstrated in animal models ^1–4^. These antibodies are effective in suppressing viremia in humans and large-scale clinical trials to test their efficacy in prevention are currently under way^2, 5–12^. However, these antibodies typically have one or more unusual characteristics including high levels of somatic hypermutation (SHM), long or very short complementarity determining regions (CDRs) and self-reactivity that interfere with their elicitation by traditional immunization.

Consistent with their atypical structural features, antibodies that broadly neutralize HIV-1 have been elicited in camelids, cows and transgenic mice with unusual pre-existing antibody repertoires^13–18^. However, even in transgenic mice that carry super-physiologic frequencies of bNAb precursors, antibody maturation required multiple immunizations with a number of different sequential immunogens. Moreover, bNAbs only developed for one of the epitopes targeted^15, 17, 18^. Consequently, elicitation of bNAbs in primates or humans remains a significant challenge.

To bypass this issue, we developed a method to reprogram mature B cells to express an anti-HIV-1 bNAb. Adoptive transfer of the engineered B cells and immunization with a single cognate antigen led to germinal center formation and antibody production at levels consistent with protection.

## Results

### Expressing antibodies in primary mature, murine B cells

To edit mature B cells efficiently, they need to be activated and cultured *in vitro*. To determine whether such cells can participate in humoral immune responses *in vivo* we used *Igh^a^* CD45.1 B cells carrying the *B1-8^hi^* heavy chain that are specific for the hapten 4-hydroxy-3-nitro-phenylacetyl (NP)^19^. B1-8^hi^ B cells were activated *in vitro* with anti-RP105 antibody for 1 - 2 days and subsequently transferred into congenically marked (*Igh^b^* CD45.2) C57BL/6J mice. Recipients immunized with NP conjugated to ovalbumin (NP-OVA) developed GCs containing large numbers of the antigen-specific, transferred B cells (Supplementary Fig. 1a,b) and produced high levels of antigen-specific IgG1 (Supplementary Fig. 1c). In addition, transfection by electroporation did not affect the ability of transferred cells to enter GCs (Supplementary Fig. 1d,e).

Despite having two alleles for each of the antibody chains, B cells express only one heavy and one light chain gene, a phenomenon referred to as allelic exclusion^20–22^. Introducing additional antibody genes would risk random combinations of heavy and light chains some of which could be self-reactive or incompatible. Thus, deletion of the endogenous chains would be desirable to prevent expression of chimeric B cell receptors (BCRs) composed of the transgene and the endogenous antibody genes. To do so, we combined endogenous Ig disruption with insertion of a transcription unit that directs expression of the heavy and light chain into the endogenous heavy chain locus. CRISPR-RNAs (crRNAs) were designed to ablate the κ–light chain because 95% of all mouse B cells express *Igk* (Fig1.a). The efficiency of k light chain deletion was measured by flow cytometry using the ratio of κ/λ cells to normalize for cell death due to BCR loss. The selected crRNAs consistently ablated Igκ expression by 70 - 80 % of B cells as measured by flow cytometry or TIDE (Tracking the Indels by DEcomposition^23^) analysis (Fig.1 b-d).

**Figure 1:**
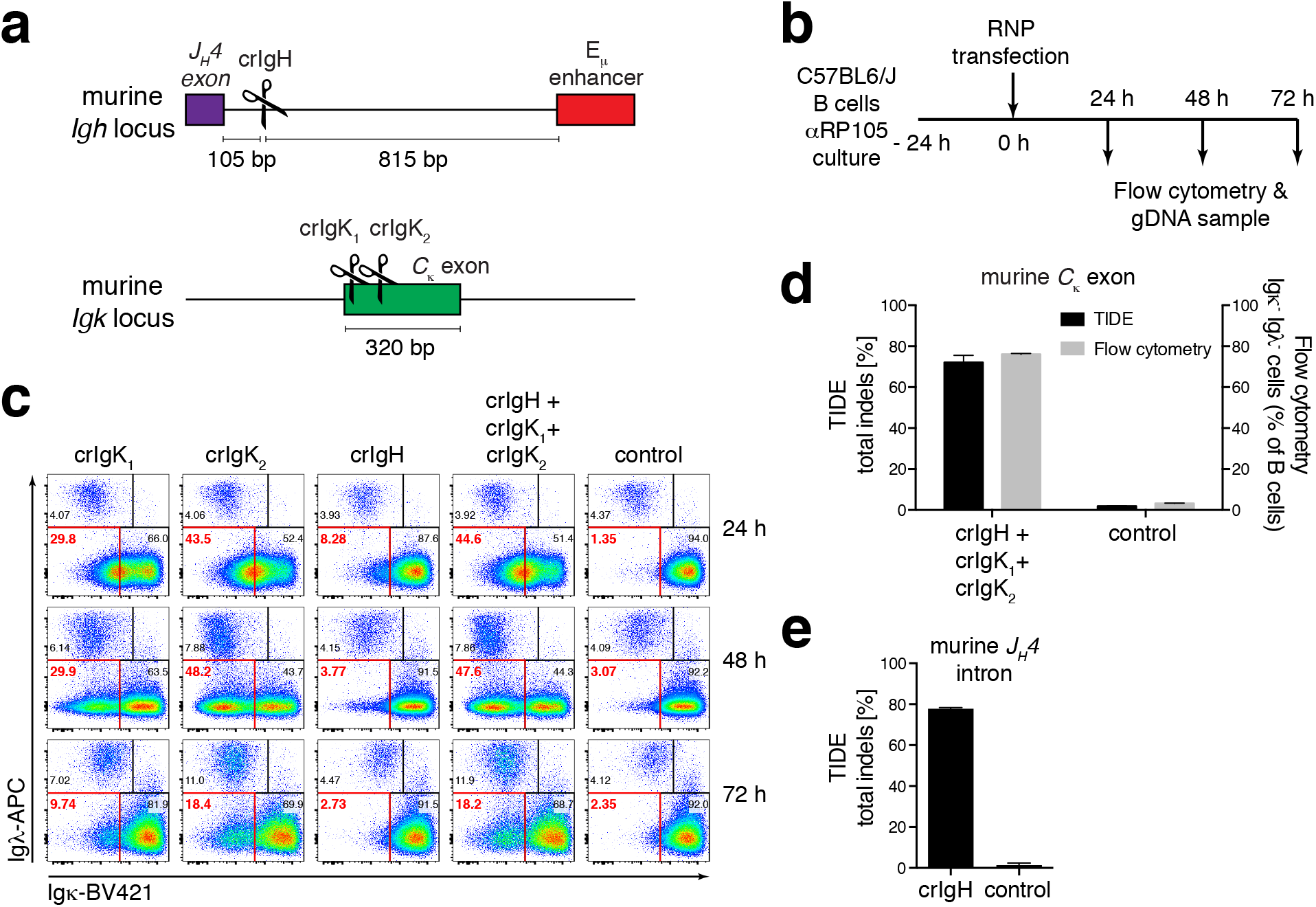
Efficient generation of indels in primary mouse B cells by CRISPR/Cas9. **(a)** Targeting scheme for *Igh* (crIgH) and *Igk* crRNA guides (crIgK_1_, crIgK_2_). **(b)** Experimental set up for (c-e). Primary mouse B cells were cultured for 24 h in the presence of anti-RP105 antibody and then transfected with Cas9 ribonucleoproteins (RNPs) and analyzed at the indicated time points. **(c)** Flow cytometric plots of cultured B cells at the indicated time points after transfection. Control uses an irrelevant crRNA targeting the HPRT gene. **(d)** Quantification of (c), percentage of Igκ^−^ Igλ^−^ B cells by flow cytometry (right y-axis) and percentage of cells containing indels in the *Igkc* exon by TIDE analysis (left y-axis). Control bars include irrelevant HPRT-targeting crRNAs or a scramble crRNA without known targets in the mouse genome. **(e)** Percentage of cells containing indels in the J_H_4 intron by TIDE analysis after targeting with crIgH or control. Bars indicate mean ± SEM in two (TIDE) or four (flow cytometry) independent experiments.

To insert a transgene into the heavy chain locus we designed crRNAs specific for the first *Igh* intron immediately 3’ of the endogenous VDJ_H_ gene segment, and 5’ of the *E_μ_* enhancer. This position was selected to favor transgene expression and allow simultaneous disruption the endogenous heavy chain (see below and ^24^). We tested 7 crRNAs and selected a high-efficiency crRNA located 110 bp downstream of the *J_H_4* intron producing 77 % indels by the TIDE assay (Fig1e, Supplementary Fig.2a,b). This location also allowed for sufficient homology to introduce a transgene, irrespective of the upstream VDJ rearrangement.

The homology-directed repair template is composed of a splice acceptor (SA) stop cassette to terminate transcription of upstream rearranged VDJ_H_, and a V_H_-gene promoter followed by cDNAs encoding *Igk*, a P2A self-cleaving sequence, and IgV_H_ with a J_H_1 splice donor (SD) site (Fig.2a). This design disrupts expression of the endogenous locus, while encoding a transcription unit directing expression of the introduced heavy and light chain under control of endogenous *Igh* gene regulatory elements. In addition, it preserves splicing of the transgenic IgV_H_ into the endogenous constant regions allowing for expression of membrane and secreted forms of the antibody as wells as different isotypes by class switch recombination. Finally, correctly targeted cells are readily identified and enumerated by flow cytometry because they bind to cognate antigen.

**Figure 2:**
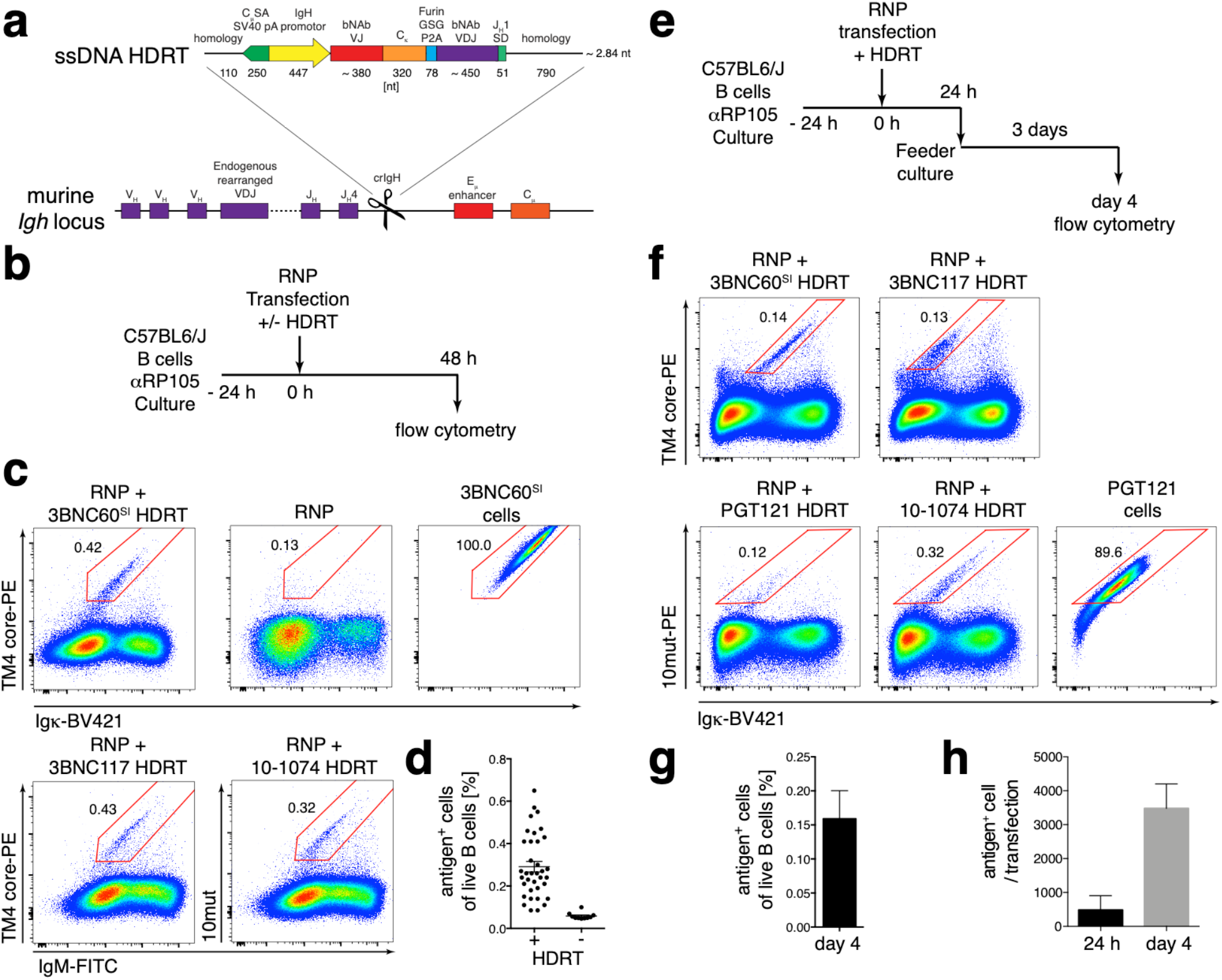
Engineering bNAb-expressing, primary, mouse B cells. **(a)** Schematic representation of the targeting strategy to create bNAb-expressing, primary mouse B cells. ssDNA homology-directed repair template (HDRT) contained 110 nt 5’ and 790 nt 3’ homology arms flanking an expression cassette. The 5’ homology arm is followed by the 111 nt long splice acceptor site and the first 2 codons of Cμ exon 1, a stop codon and a SV40 polyadenylation signal (CμSA SV40 pA). Then the mouse *Ighv4-9* gene promoter, the leader, variable and joining regions (VJ) of the respective antibody light chain and mouse k constant region (C_κ_) are followed by a furin-cleavage site, a GSG-linker and a P2A self-cleaving oligopeptide sequence (P2A), the leader, variable, diversity and joining regions (VDJ) of the respective antibody heavy chain and 45 nt of the mouse J_H_1 intron splice donor site to splice into downstream constant regions. **(b)** Experimental setup for (c). **(c)** Flow cytometric plots of primary, mouse B cells, activated and transfected with RNPs targeting the *Igh* J_H_4 intron and *Igkc* exon with or without ssDNA HDRTs encoding the 3BNC60^SI^, 3BNC117 or 10-1074 antibody. Non-transfected, antigen-binding B cells from 3BNC60^SI^ knock-in mice cultured the same way are used as control for gating. **(d)** Quantification of (c). Each dot represents one transfection. Data from 7 independent experiments. **(e)** Experimental set up for (f-h) **(f)** Flow cytometric plots of primary, mouse B cells, activated and transfected using ssDNA HDRT encoding the antibodies 3BNC60^SI^, 3BNC117, PGT121 or 10-1074. B cells were expanded on feeder cells for 3 days. Cultured, non-transfected, antigen-binding B cells from PGT 121 knock-in mice are shown for gating. **(g)** Quantification of (f). **(h)** Total number of antigen binding B cells before (24 h) or after 3 days (day 4) of feeder culture. Bars indicate mean ± SEM of combined data from 2 independent experiments.

A number of methods for producing ssDNA homology directed repair templates (HDRTs) were compared. The most reproducible and least cytotoxic involved digestion of plasmids with sequence-specific nickases, and ssDNA purification by agarose gel electrophoresis (Supplementary Fig. 3a-c,^25, 26^).

Co-transfection of the ssDNA template with pre-assembled Cas9 ribonucleoproteins (RNPs) containing the crRNAs resulted in expression of the encoded anti-HIV antibody in 0.1 - 0.4 % of mouse B cells by antigen-specific flow cytometry (TM4 core^27, 28^ or 10mut^29^) (Fig.2c,d). Transgene expression was stable over the entire culture period of 3 days on feeder cells^30^, during which the overall number of B cells expanded by 6 to 20-fold (Fig.2e-h). However, expression of transgenic antibodies differed depending on the antibody and were generally reflective of their expression in knock-in mouse models (Fig.2c, f)^16, 18, 28, 29, 31^.

To determine whether edited cells are allelically excluded we transfected *Igh^a/b^* B cells with 3BNC60^SI^, a chimeric antibody composed of the mature heavy chain and germline light chain of the anti-HIV bNAb 3BNC60 (Supplementary Fig.4b, c). The majority of edited cells expressing the 3BNC60^SI^ transgene, expressed it using either *Igm^a^* or *Igm^b^* allele as determined by flow cytometry. Only 5.21 % of 3BNC60^SI^-expressing B cells showed co-expression of both IgM^a^ and IgM^b^ indicative of allelic inclusion of the endogenous allele or successful integration of the transgene into both alleles. Thus, the majority of edited B cells express only the transgene.

We conclude that mature mouse B cells can be edited *in vitro* to produce anti-HIV-1 bNAbs from the *Igh* locus.

### Antibody gene editing in human B cells

To determine whether this method could be adapted to edit human B cells we isolated them from peripheral blood of healthy volunteers and activated them using an anti-human RP105 antibody^32^. Analogous crRNAs were selected for targeting the human *IGKC* and the first intron 3’ of *IGHJ6* (Fig.3a-d, Supplementary Fig.5a-b). The best IGKC-targeting crRNA caused 85 % of κ-bearing B cells to lose BCR expression, whereas λ-bearing cells increased proportionally indicating that they were unaffected. TIDE analysis of the J_H_6 intron sequences showed that the most efficient crRNA induced 64% indels. In conclusion, activation of human, primary B cells with anti-RP105 allows efficient generation of indels using Cas9 RNPs.

**Figure 3:**
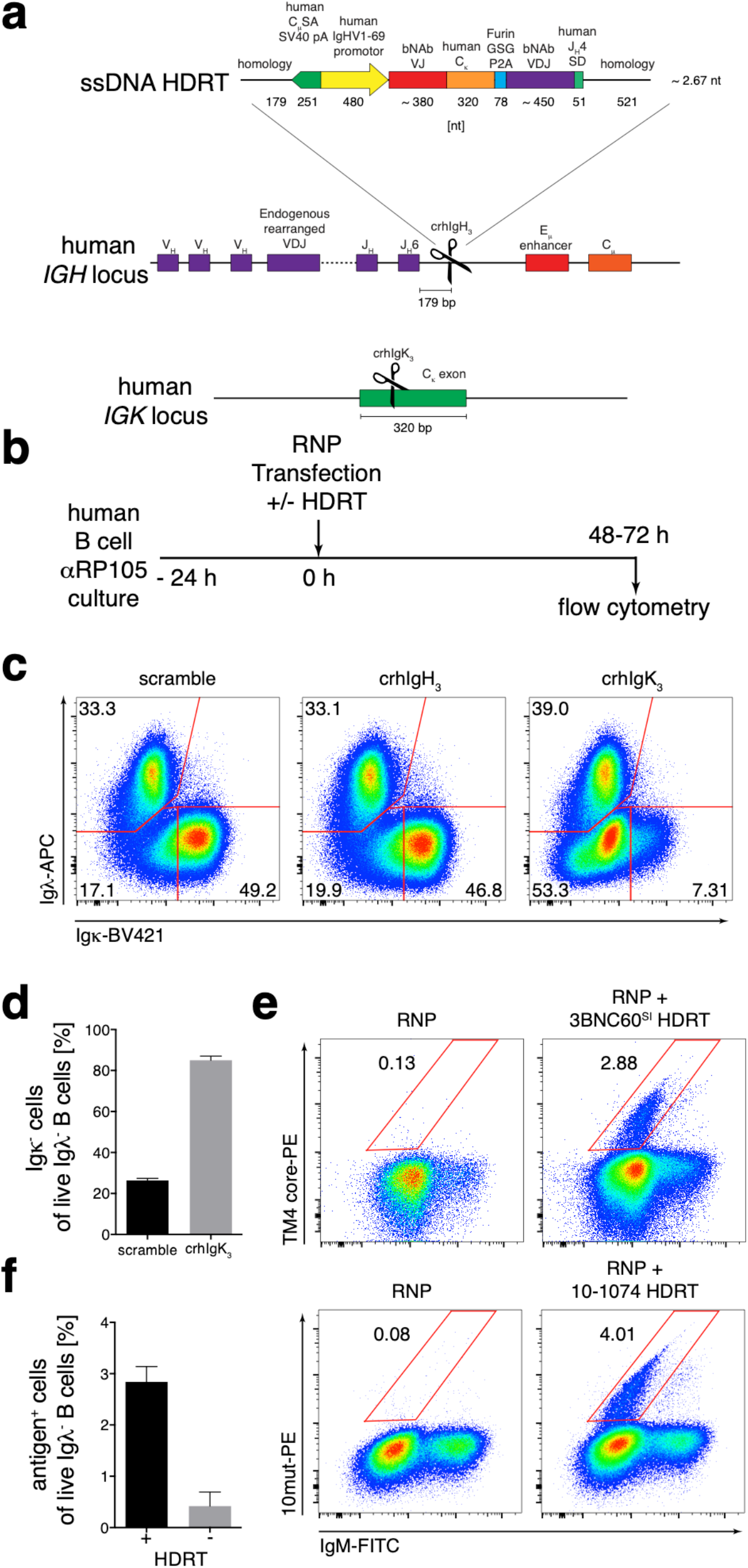
Engineering bNAb-expressing, primary, human B cells. **(a)** Schematic representation of the targeting strategy to create bNAb-expressing, primary human B cells. The ssDNA HDRT is flanked by 179 nt and a 521 nt homology arms. The central expression cassette contains 112 nt of the human splice acceptor site and the first 2 codons of Cμ exon 1, a stop codon and a SV40 polyadenylation signal (CμSA SV40 pA). Then the human *IGHV1-69* gene promoter, the leader, variable and joining regions (VJ) of the respective antibody light chain and human k constant region (Cĸ) are followed by a furin-cleavage site, a GSG-linker and a P2A self-cleaving oligopeptide sequence (P2A), the leader, variable, diversity and joining regions (VDJ) of the respective antibody heavy chain and 50 nt of the human J_H_4 intron splice donor site to splice into downstream constant regions. **(b)** Experimental set up for c,d. Primary human B cells were cultured for 24 h in the presence of anti-RP105 antibody and then transfected with RNPs ± HDRT. **(c)** Flow cytometric plots of primary human B cells 48 h after transfection with RNPs containing crRNAs without target (scramble) or targeting the *IGHJ6* intron or the *IGKC* exon. **(d)** Quantification of (c). Combined data from 3 independent experiments is shown. **(e)** Flow cytometric plots of antigen binding by Igλ^−^ primary human B cells 72 h after transfection of RNPs targeting both the *IGHJ6* intron and the *IGKC* exon with or without HDRTs encoding 3BNC60^SI^ or 10-1074. **(f)** Quantification of (e). Bars indicate mean ± SEM of combined data from 2 independent experiments with 2 - 4 replicates each.

To target bNAbs into the human J_H_6 intron we adapted the ssDNA HDRT and replaced mouse with human homology arms, the human Cμ splice acceptor, the human *IGHV1-69* promoter, a codon-modified human *IGKC* constant region to avoid targeting by crRNAs and the human *J_H_4* splice donor (Fig 3a). In contrast to mouse cells, 2.9 – 4 % of λ^−^ B cells expressed 3BNC60^SI^ or 10-1074 antibodies respectively as determined by flow cytometry using the cognate antigen (Fig 3e,f). Thus, the efficiency of transgene integration is at least 10-times higher in human B cells. Furthermore, viability was also higher in human B cells, ranging from 60 to 85 % of live cells after transfection (Supplementary Fig.5c).

We conclude that primary human B cells can be edited by CRISPR/Cas9 to express anti-HIV bNAbs, and that this is significantly more efficient than in mouse B cells.

### Adoptive transfer of antibody-edited B cells

To determine whether edited B cells can participate in immune responses, we adoptively transferred mouse 3BNC60^SI^-edited *Igh^b^* B cells, into congenically-marked *Igh^a^* wild type mice and immunized with the high-affinity, cognate antigen TM4 core in Ribi adjuvant (Fig.4a). Transgene-specific responses were detected using anti-idiotypic antibodies as an initial capture reagent in ELISA. Similar to endogenous humoral immune responses, transgenic antibodies were detected on day 7 after immunization, they peaked at day 14 and started to decrease by day 21 (Fig.4b, c). Importantly, the transgenic immune response included secondary isotypes indicating that the re-engineered locus supports class switch recombination (Fig.4c). Finally, the magnitude of the response was directly correlated to the number of transferred cells. However, prolonged *in vitro* culture under the conditions tested decreased the efficiency of antibody production *in vivo* (Fig.4d).

**Figure 4:**
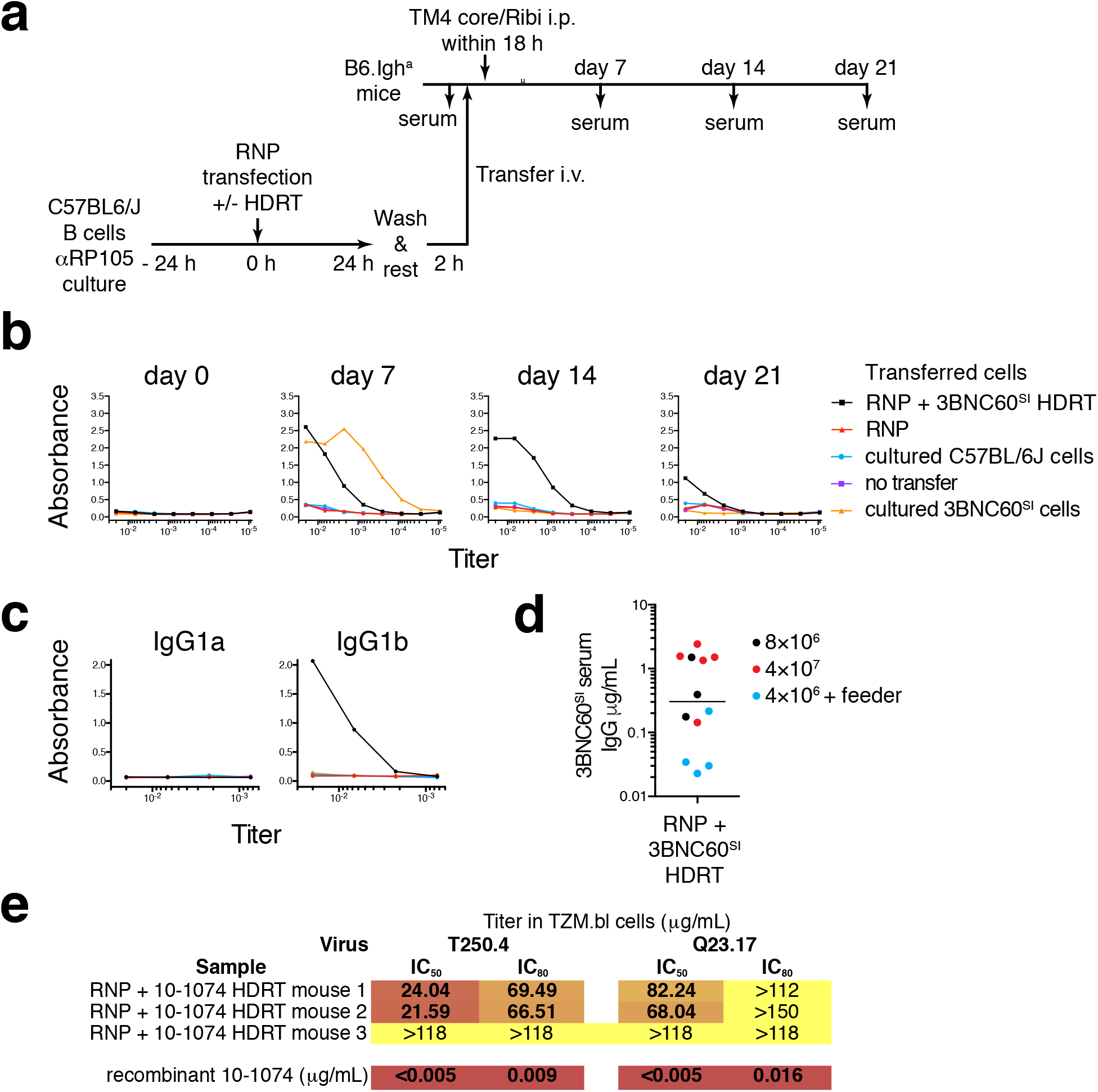
Engineered bNAb-expressing, primary, mouse B cells participate in humoral immune responses *in vivo*. **(a)** Experimental set up for (b-e). i.v. intravenous; i.p. intraperitoneal. **(b)** Anti-3BNC60^SI^ idiotype-coated, mouse IgG ELISA of sera from mice adoptively transferred with the indicated B cells and immunized with the cognate antigen TM4 core at the indicated time points. Representative plots of 7 independent experiments. **(c)** Anti-3BNC60^SI^ idiotype-coated, mouse IgG1^a^ or IgG1^b^ ELISA of day 14 sera, as above. Representative plots of 2 independent experiments. **(d)** 3BNC60^SI^ serum IgG levels 14 days after immunization in mice transferred with 3BNC60^SI^-edited cells, numbers of total B cells/mouse at transfection are indicated. Cells were transferred either 24 h after transfection or after additional culture on feeder cells as in Fig.2d. Determined by anti-3BNC60^SI^ idiotype-coated, mouse IgG ELISA over 7 independent experiments. Each dot represents one mouse. **(e)** TZM.bl neutralization data of protein G-purified serum immunoglobulin days 14 – 21 after immunization from mice treated as in (a) but transfected with 10-1074 HDRT and immunized with cognate antigen 10mut. Combined data from 2 independent experiments are shown.

To determine whether the transferred cells retained the ability to produce neutralizing antibodies we used B cells that were edited to produce 10-1074, a potent bNAb, or 3BNC60^SI^ a chimeric antibody with limited neutralizing activity^31, 33^. 4 × 10^7^ transfected B cells were transferred into wild type Igh^a^ mice that were subsequently immunized with the appropriate cognate antigen 10mut^29^ or TM4 core^16, 27, 28, 31^. Serum IgG was purified from 3 mice each that received an estimated ~10^3^ edited B cells expressing 10-1074 or 3BNC60^SI^. The serum antibodies were tested for neutralizing activity in the TZM-bl assay^34^. Two of the 3 mice that received 10-1074 edited cells showed IC50s of 21.59 μg/mL and a third reached 49 % neutralization at 118 μg/mL (corresponding to approximately 1:500 and 1:100 dilution of serum, Fig.4e, Supplementary Fig.6a, b). As expected, neutralizing activity was not detected in mice receiving 3BNC60^SI^ because this antibody is 2 - 3 orders of magnitude less potent against the tested viral strains than 10-1074 (Supplementary Fig.6e).

We conclude that edited B cells can be recruited into immune responses and produce sufficient antibody to confer potentially protective levels of humoral immunity^35^.

## Discussion

T cells can be reprogrammed to express specific receptors using retrogenic methods^36–38^ or non-viral CRISPR/Cas9 genome targeting^26^. In contrast, B cell receptor reprogramming in primary cells using retroviruses has not been successful^39^. Moreover, although antibody heavy chains have been targeted into human B cells using CRISPR/Cas9^40^, little is known about how CRISPR/Cas9 genome targeting might be used to introduce complete antibody genes into mature B cells that retain the ability to participate in immune responses *in vivo*.

We have developed a method to produce transgenic antibodies in primary mouse and human B cells using CRISPR/Cas9. The new method involves short term culture *in vitro*, silencing of the endogenous *Ig* genes, and insertion of a bi-cistronic cDNA into the *Igh* locus. Mouse B cells edited to express an anti-HIV-1 bNAb by this method can produce transgenic antibody levels that are protective in animal models^35, 41–43^.

Mouse and human B lymphocytes typically express a single antibody despite having the potential to express 2 different heavy chains and 4 different light chains. Theoretically the combination could produce 8 different antibodies and a series of additional chimeras that could interfere with the efficiency of humoral immunity and lead to unwanted autoimmunity. Allelic exclusion prevents this from happening and would need to be maintained by any gene replacement strategy used to edit B lymphocytes. In addition, genetic editing is accompanied by safety concerns due to off-target double strand breaks and integrations.

The approach reported maintains allelic exclusion in part by ablating the *Igkc* gene. In the mouse, 95% of B cells express *Igkc*. In the absence of *Igkc* expression these cells will die by apoptosis because they cannot survive unless they continue to express a B cell receptor^44, 45^. Since the introduction of the transgene into the heavy chain locus disrupts endogenous *Igh* expression, editing maintains allelic exclusion because only cells expressing the introduced antibody can survive. Our strategy also interferes with the survival of cells that suffer off-target integration events, because such cells would be unable to express the B cell receptor and they too would die by apoptosis.

A potential issue is that there are two heavy chain alleles in every B cell and allelic exclusion would be disrupted if the transgene were only integrated in the non-productive *Igh* allele allowing for expression of the original productive *Igh*. However, our flow cytometry data indicates that this is a very rare event. Thus, either both alleles are targeted or the occasional remaining endogenous *Igh* gene is unable to pair with the transgenic *Igk*

In contrast to the mouse, *IGL* is expressed by 45 % of all B cells in humans. Therefore, this locus would either need to be ablated, or alternatively, cells expressing *IGL* could be removed from the transferred population by any one of a number of methods of negative selection.

Similar to antibody transgenes in mice, expression of the edited BCR varied between different antibodies. Some combinations of heavy and light chains were refractory to expression in mature B cells. In addition, although the level of B cell receptor expression was within the normal range, it was generally in the low end compared to polyclonal B cells. This is consistent with generally lower level expression of a similar transgene in knock-in mice^24^. Low BCR expression could also be due to the bi-cistronic design since expression was higher in knock-in mice that expressed the identical Ig from the native *Igk* and *Igh* loci^31^. Nevertheless, expression levels were adequate to drive antigen-induced antibody production *in vivo*.

bNAb mediated protection against infection with simian-human immunodeficiency viruses in macaques requires IC50 neutralizing titers of 1:100^35, 41–43^. Thus the titers achieved by CRISPR/Cas9 edited B cells in mice would be protective if they could be translated to macaques and by inference humans. Moreover, our neutralization measurements may be an underestimate since we excluded bNAbs produced as IgM or isotypes other than IgG.

CAR T cell therapy typically involves transfer of millions of edited cells to achieve a therapeutic effect. Whether similar numbers of edited B cells would also be required to achieve protective levels of humoral immunity can only be determined by further experimentation in primate models. In addition, the longevity of the antibody response produced by edited B cells, and its optimization by boosting or adjuvant choice will require further experimentation.

Most protective vaccine responses depend on humoral immunity. Neutralizing antibody responses are readily elicited for most human pathogens, but in some cases, including HIV-1, it has not yet been possible to do so. The alternatives include passive antibody infusion, which has been an effective means of protection since it was discovered at the turn of the last century. We have shown that passive transfer of mouse B cells edited by CRISPR/Cas9 can also produce protective antibody levels *in vivo*. This proof of concept study demonstrates that humoral immune responses can be engineered by CRISPR/Cas9. The approach is not limited to HIV-1 and can be applied to any disease requiring a specific antibody response.

## Methods

### crRNA design

crRNAs were designed with the MIT guide design tool (crispr.mit.edu), CHOPCHOP (chopchop.cbu.uib.no) and the IDT crRNA design tool (www.idtdna.com). Designs were synthesized by IDT as Alt-R CRISPR-Cas9 crRNAs. crRNA sequences are listed in Supplementary Table 1.

### ssDNA HDRT preparation

HDRT sequences, listed in Supplementary Table 2, were synthesized as gBlocks (IDT) and cloned using *NheI* and *XhoI* (NEB) into vector pLSODN-4D from the long ssDNA preparation kit (BioDynamics Laboratories, Cat.# DS620). ssDNA was prepared following the manufacturer’s instructions with the following modifications: In brief, 2.4 mg sequence verified vector was digested at 2 μg/μL in NEB 3.1 buffer with 1200 U Nt.*BspQI* for 1 h at 50 °C followed by addition of 2400 U *XhoI* (NEB) and incubation for 1 h at 37 °C. Digests were desalted by ethanol precipitation and resuspended in water at < 1 μg/μL. An equal volume of formamide gel-loading buffer (95 % de-ionized formamide, 0.025 % bromophenol blue, 0.025 % xylene cyanol, 0.025 % SDS, 18 mM EDTA) was added and heated to 70 °C for 5 min to denature dsDNA. Denatured samples were immediately loaded into dye-free 1 % agarose gels in TAE and run at 100 V for 3 h. Correctly sized bands were identified by partial post-stain with GelRed (Biotium), then excised and column purified (Machery Nagel Cat.# 740610.20 or 740609.250) according to the manufacturer’s instructions. Eluate was ethanol precipitated, resuspended in water, adjusted to 2.5 μg/μL and stored at −20 °C.

### Murine cell culture

Mature, resting B cells were obtained from mouse spleens by forcing tissue through a 70 μm mesh into PBS containing 2 % heat-inactivated fetal bovine serum (FBS). After ACK lysis for 3 min, untouched B cells were enriched using anti-CD43 magnetic beads (MACS) according to manufacturer’s protocol (Miltenyi Biotec) obtaining > 95 % purity. 3.2 × 10^7^ cells/10 cm dish (Gibco) were cultured at 37 °C 5 % CO_2_ in 10 mL mouse B cell medium consisting of RPMI-1640, supplemented with 10 % heat-inactivated FBS, 10 mM HEPES, antibiotic-antimycotic (1X), 1 mM sodium pyruvate, 2 mM L-glutamine and 53 μM 2-mercaptoethanol (all from Gibco) and activated with 2 μg/mL anti-mouse RP105 clone RP/14 (produced in house or BD Pharmingen Cat.# 562191).

NB-21 feeder cells^30^ were maintained in DMEM supplemented with 10 % heat-inactivated FBS and antibiotic-antimycotic (1X). For co-culture, feeder cells were irradiated with 80 Gy and seeded simultaneously with B cells, 24 h after transfection, into B cell culture medium supplemented with 1 ng/mL recombinant mouse IL-4 (PeproTech Ca.# 214-14) and 2 μg/mL anti-mouse RP105 clone RP/14.

### Human cell culture

Frozen human leukapheresis samples were collected after signed informed consent in accordance with Institutional Review Board (IRB) approved protocol TSC-0910. Cells were thawed in a 37°C water bath and resuspended in human B cell medium composed of RPMI-1640, supplemented with 10 % heat-inactivated FBS or human serum, 10 mM HEPES, antibiotic-antimycotic (1X), 1 mM sodium pyruvate, 2 mM L-glutamine and 53 μM 2-mercaptoethanol (all from Gibco). B cells were isolated using EasySep human naïve B cell Enrichment Kit (Stemcell Cat.# 19254) according to the manufacturer’s instructions and cultured in the above medium supplemented with 2 μg/mL anti-human RP105 antibody clone MHR73-11 (BioLegend Cat.# 312907).

### RNP preparation and transfection

Per 100 μL transfection, 1 μL of 200 μM crRNA and 1 μL 200 μM tracrRNA in duplex buffer (all IDT) were mixed, denatured at 95 °C for 5 min, re-natured for 5 min at room temperature. 5.6 μL PBS and 2.4 μL 61 μM Cas9 V3 (IDT, Cat.# 1081059) were added and incubated for 15 - 30 min. If required RNPs were mixed at the following ratios: 50 % crIgH, 25 % crIgK1 and 25 % crIgK2 (mouse) or 50% crhIgH3 and 50 % crhIgK3 (human). 4 μL 100 μM electroporation enhancer in duplex buffer or 4 μL HDRT at 2.5 μg/μL were added to 10 μL mixed RNP and incubated for a further 1-2 min.

24 h after stimulation, activated mouse or human B cells were harvested, washed once in PBS and resuspended in Mouse B cell Nucleofector Solution with Supplement (murine B cells) or Primary Cell Nucleofector Solution 3 with Supplement (human B cells) prepared to the manufacturer’s instructions (Lonza) at a concentration of 4 - 5 × 10^6^ cells / 86 μL. 86 μL cells were added to the RNP/HPRT mix, gently mixed by pipetting and transferred into nucleofection cuvettes and electroporated using an Amaxa IIb machine setting Z-001 (murine B cells) or Amaxa 4D machine setting EH-140 (human B cells). Cells were immediately transferred into 6-well dishes containing 5 mL prewarmed mouse or human B cell medium supplemented with the relevant anti-RP105 antibody at 2 μg/mL and incubated at 37°C 5 % CO_2_ for 24 h before further processing.

### TIDE assay

Genomic DNA was extracted from 0.5 - 5 × 10^5^ cells by standard phenol/chloroform extraction 24 - 42 h after transfection. PCRs to amplify human or mouse Ig loci targeted by CRISPR-Cas9 were performed using Phusion Green Hot Start II High-Fidelity polymerase (Thermo Fisher Cat.# F-537L) and primers listed in Supplementary Table 3. Thermocycler was set to 40 cycles, annealing at 65°C for 30 s and extending at 72°C for 30s. PCR product size was verified by gel electrophoresis, bands gel-extracted and sent for Sanger sequencing (Genewiz) using the relevant PCR primers. ab1 files were analyzed using the TIDE web tool (tide.nki.nl) using samples receiving scamble or irrelevant HPRT-targeting crRNA as reference^23^.

### Flow cytometry

Mouse spleens were forced through a 70 μm mesh into FACS buffer (PBS containing 2 % heat-inactivated FBS and 2 mM EDTA) and red blood cells were lysed in ACK lysing buffer (Gibco) for 3 min. Cultured cells were harvested by centrifugation. Then cells were washed and Fc-receptors blocked for 15 min on ice. Cells were stained for 20 min on ice with antibodies or reagents listed in Supplementary table 4 and depending on the stain, washed again and secondary stained for another 20 min on ice before acquisition on a BD LSRFortessa. GC B cells were gated as single/live, B220^+^, CD38^−^ FAS^+^, GL7^+^, IgD^−^. Allotypic markers CD45.1 and CD45.2 were used to track adoptively transferred B cells.

### Mice

C57BL/6J and B6.Igh^a^ (B6.Cg-Gpi1^a^ Thy1^a^ Igh^a^/J) and B6.SJL were obtained from the Jackson Laboratory. Igh^a/b^ mice were obtained by intercrossing B6.Igh^a^ and B6.SJL mice. B1-8^hi 19^, 3BNC60^SI 31^ and PGT121^18, 29^ strains were generated and maintained in our laboratory. Experiments used age and sex-matched animals. All experiments were performed with authorization from the Institutional Review Board and the Rockefeller University IACUC.

### Cell transfers and immunizations

After culture, mouse B cells were harvested at the indicated time points and resuspended in mouse B cell medium without anti-RP105 antibody and rested for 2 - 3 h at 37°C, 5 % CO_2_. Then cells were washed once in PBS and resuspended in 200 μL PBS/mouse containing the indicated number of initially transfected cells. 200 μL cell suspension/mouse were injected intravenously via the retroorbital sinus. Number of transferred, edited B cells was estimated as follows: Number of cells transfected × 20 % survival x 0.15 - 0.4 % transfection efficiency x 50 % handling/proliferation x 5 % transfer efficiency^31^. Mice were immunized intraperitoneally within 24 h after cell transfer with 200 μL containing 10 μg TM4 core^27^ or 10mut^29^ in PBS with 50 % Ribi (Sigma Adjuvant system, Sigma Aldrich) prepared to the manufacturer’s instructions. Mice were bled at the indicated time points from the submandibular vein. Blood was allowed to clot and then serum was separated by centrifugation for 10 min at 20817 g. Serum was stored at −20°C.

### Anti-idiotypic antibody

IgG producing hybridomas were isolated from mice immunized with iGL-VRC01 at the Frederick Hutchinson Cancer Research Center Antibody Technology Resource. Hybridoma supernatants were screened against a matrix of inferred germline (iGL) VRC01 class antibodies as well as irrelevant iGL-antibodies using a high throughput bead-based assay. One anti-idiotypic antibody, clone iv8, bound to additional VRC01 class antibodies, but it also bound to a chimeric antibody with an iGL-VRC01 class light chain paired with the 8ANC131 heavy chain (which is derived from VH1-46), and to 3BNC60^SI^.

### ELISAs

For determination of 3BNC60^SI^ levels, Corning 3690 half-well 96-well plates were coated overnight at 4°C with 25 μL/well of 2 μg/mL human anti-3BNC60^SI^ (clone iv8) IgG in PBS, then blocked with 150 μL/well PBS 5% skimmed milk for 2 h at room temperature (RT). Sera were diluted to 1:50 with PBS and 7 subsequent 3-fold dilutions. Recombinant 3BNC60^SI^ (produced in house as mouse IgG1,k) was diluted to 10 μg/mL in PBS followed by six 5-fold dilutions. Blocked plates were washed 4-times with PBS 0.05 % Tween 20 and incubated with 25 μL diluted sera or antibody for 2 h at RT. Binding was revealed by either anti-mouse IgG-horseradish peroxidase (HRP) (Jackson ImmunoResearch, Cat.# 115-035-071) or anti-mouse IgG1a-biotin (BD Pharmingen Cat.# 553500) or anti-mouse IgG1b-biotin (BD Pharmingen Cat.# 553533), all diluted 1:5000 in PBS, 25 μL/well and incubation for 1 h at RT. Biotinylated antibodies were subsequently incubated with Streptavidin-HRP (BD Pharmingen Cat.# 554066), diluted 1:1000 in PBS, 25 μL/well for 30 min at RT. Plates were washed 4-times with PBS 0.05 % Tween 20 in between steps and 6 times before addition of substrate using a Tecan Hydrospeed microplate washer. HRP activity was determined using TMB as substrate (Thermo Scientific Cat.# 34021), adding 50 μL/well. Reactions were stopped with 50 μL/well 2 M H2SO4 and read at 450 and 570 nm on a FLUOstar Omega microplate reader (BMG Labtech). Data were analyzed with Microsoft Excel and GraphPad Prism 6.0. Absolute 3BNC60^SI^ titers were interpolated from sigmoidal fits of recombinant 3BNC60^SI^ standard curves.

For determination of NP-binding antibodies the following modifications applied. Plates were coated with 10 μg/mL NP31-bovine serum albumin (BSA, Biosearch Technologies) and blocked with PBS 3 % BSA. Sera, antibodies and secondary reagents were diluted in PBS 1% BSA 0.05 % Tween20.

### Neutralization assays

Collected mouse serum was pooled and IgG purified using protein G Ab SpinTraps (GE Healthcare Cat.# 28-4083-47) then concentrated and buffer-exchanged into PBS using Amicon Ultra 30K centrifugal filter units (Merck Millipore Cat.# UFC503024) according to the manufacturers’ instructions.

TZM-bl assays were performed as previously described^34^. Neutralizing activity was calculated as a function of the reduction in Tat-inducible luciferase expression in the TZM-bl reporter cell line in a single round of virus infection.

## Acknowledgements

We would like to thank H.B. Gristick, J.R. Keeffe and P.J. Bjorkman for providing 10mut protein and G. Kelsoe for providing NB-21 feeder cells. S. Tittley and T. Eisenreich for help with mouse colony management, T. Hägglöf and K. Yao for technical help, P. C. Rommel, E.E. Kara, E.S. Thientosapol, A.P. West Jr., G.D. Victora and all members of the Nussenzweig laboratory for discussion. This work was supported by the Bill and Melinda Gates Foundation Collaboration for AIDS Vaccine Discovery (CAVD) grants OPP1092074, OPP1124068, OPP1194977 (M.C.N.) and OPP1146996 (M.S.S.); NIH grants 1UM1 AI100663 and R01AI-129795 (M.C.N.); and the Robertson fund. H.H. was supported by the Cancer Research Institute Irvington Fellowship. M.C.N. is a Howard Hughes Medical Institute Investigator.

## Author contributions

H.H. and M.J. conceived, planned and performed experiments, analyzed data, and wrote the manuscript. M.H. performed experiments and prepared ssDNA HDRT. P.D. assisted with experimental design and ELISAs. D.Y. and A.G. expressed antibodies. A.T.G., J.J.T. and L.S. designed and provided iv8 antibody. M.S.S. performed neutralization assays. M.C.N. planned and supervised experiments, analyzed data, and wrote the manuscript.

