## Supplementary Figures for "Antigen-specific humoral immune responses by CRISPR/Cas9-edited B cells"

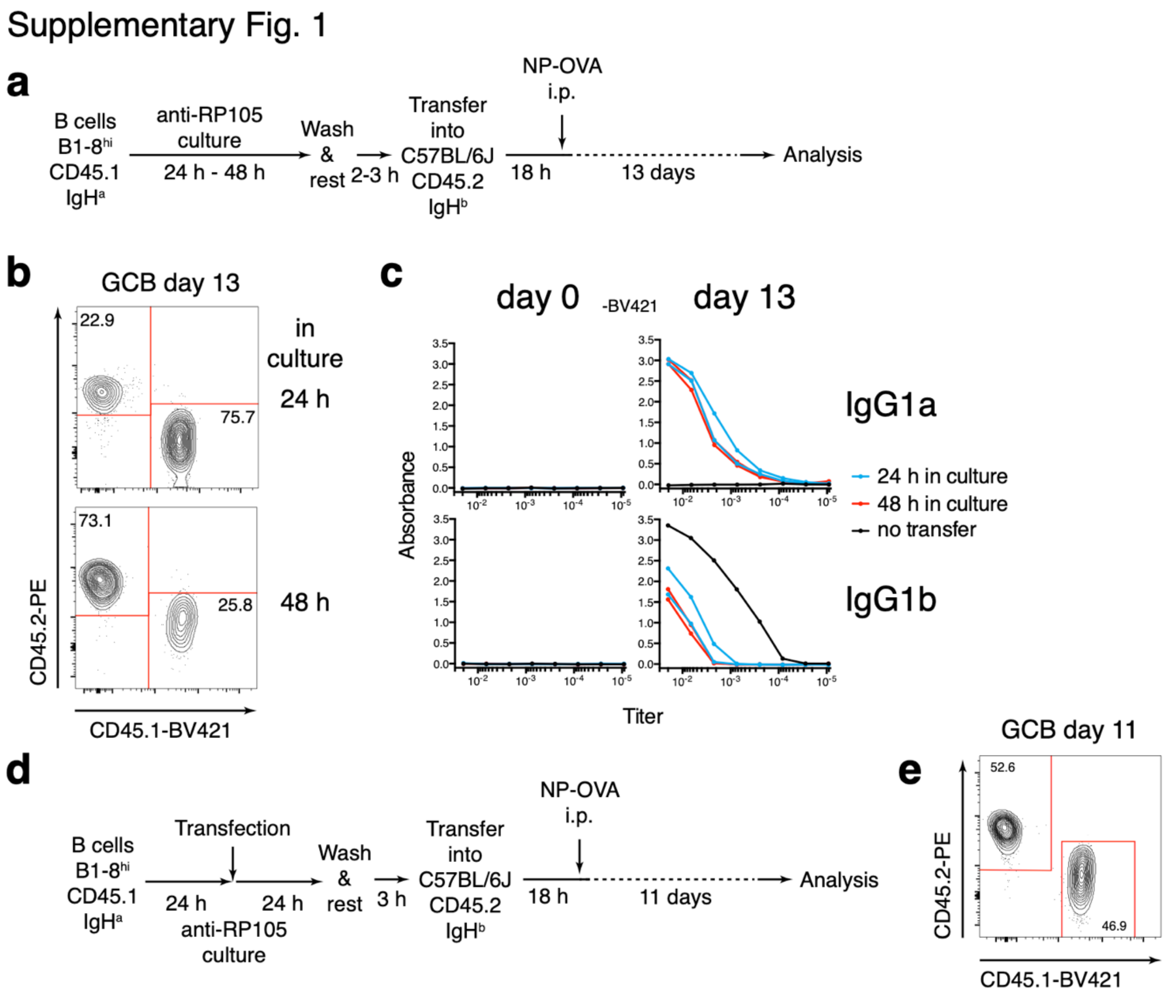


**Supplementary Figure 1: Cultured B cells participate in humoral immune responses.**

**(a)** Schematic representation of the experimental set up for (b) and (c). B1-8$^{hi}$ CD45.1 Igh$^{a}$ cells were cultured for 24 or 48 h in the presence of anti-RP105 antibody, then rested for 2 - 3 h without antibody and then transferred into C57BL6/J (CD45.2 Igh$^{b}$) recipients. 18 h later, mice were immunized with NP-OVA i.p. and mice were analyzed 2 weeks later. **(b)** Flow cytometric plots gated on CD38$^{-}$Fas$^{+}$GL7$^{+}$IgD$^{-}$ GC B cells 2 weeks after transfer. **(c)** Pre-immune (day 0) and day 13 ELISA titers of anti-NP IgG1$^{a}$ or IgG1$^{b}$. **(d)** Schematic representation of the experimental set up for (e). B1-8$^{hi}$ CD45.1

$Igh^a$ cells were cultured for 24 h and transfected with plasmid DNA. 24 h after transfection cells were transferred and analyzed as in (a). **(e)** Flow cytometric plots gated on $CD38^{-}Fas^{+}GL7^{+}IgD^{-}$ GC B cells 11 days after transfer. Data are representative of 2-3 independent experiments.

#### Supplementary Fig. 2

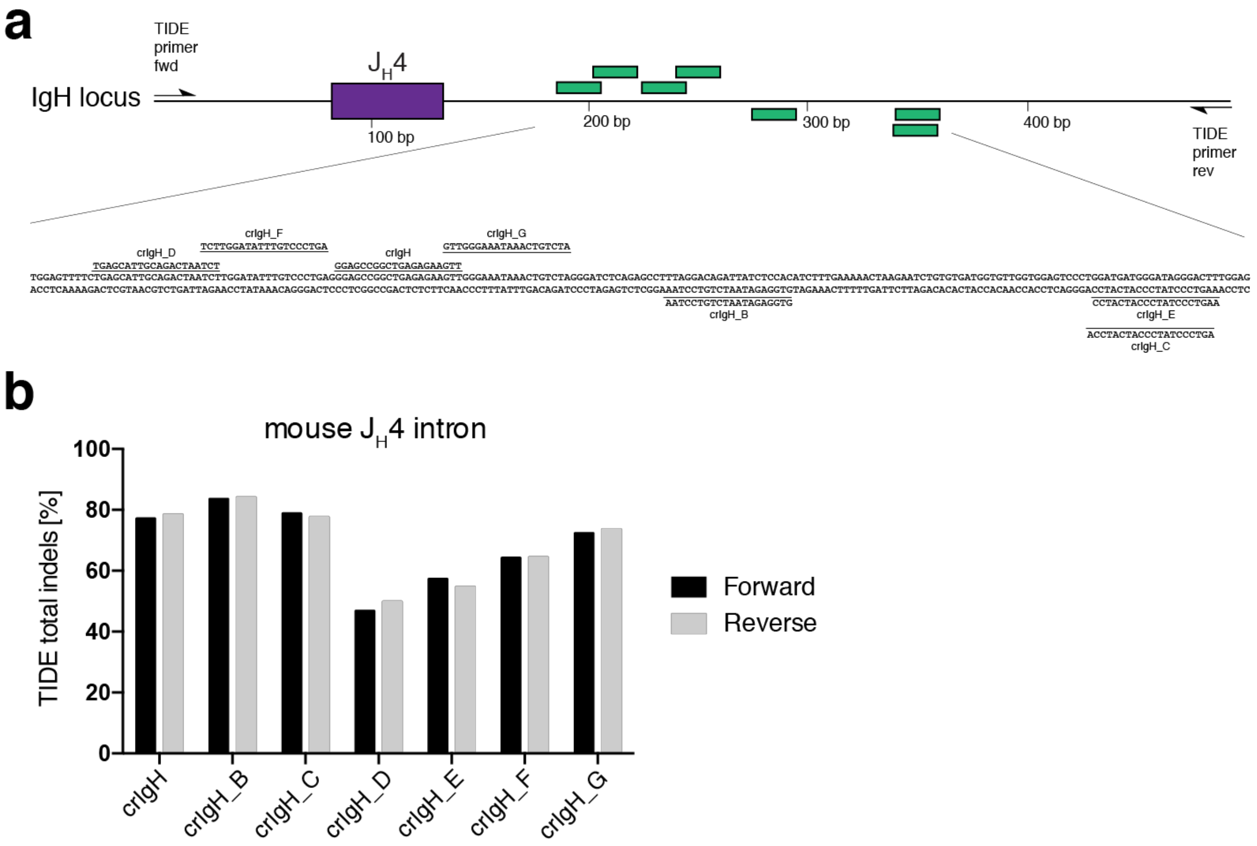


**Supplementary Figure 2: Identifying an optimal mouse *Igh* crRNA.**

**(a)** Schematic representation of the mouse *Igh* locus around $J_H4$. Location and sequence of tested guide RNAs is indicated below. **(b)** TIDE assay comparing the efficiency of creating indels of the crRNAs indicated in (a). Forward/reverse indicate sequencing with forward/reverse primers respectively.

Supplementary Fig. 3

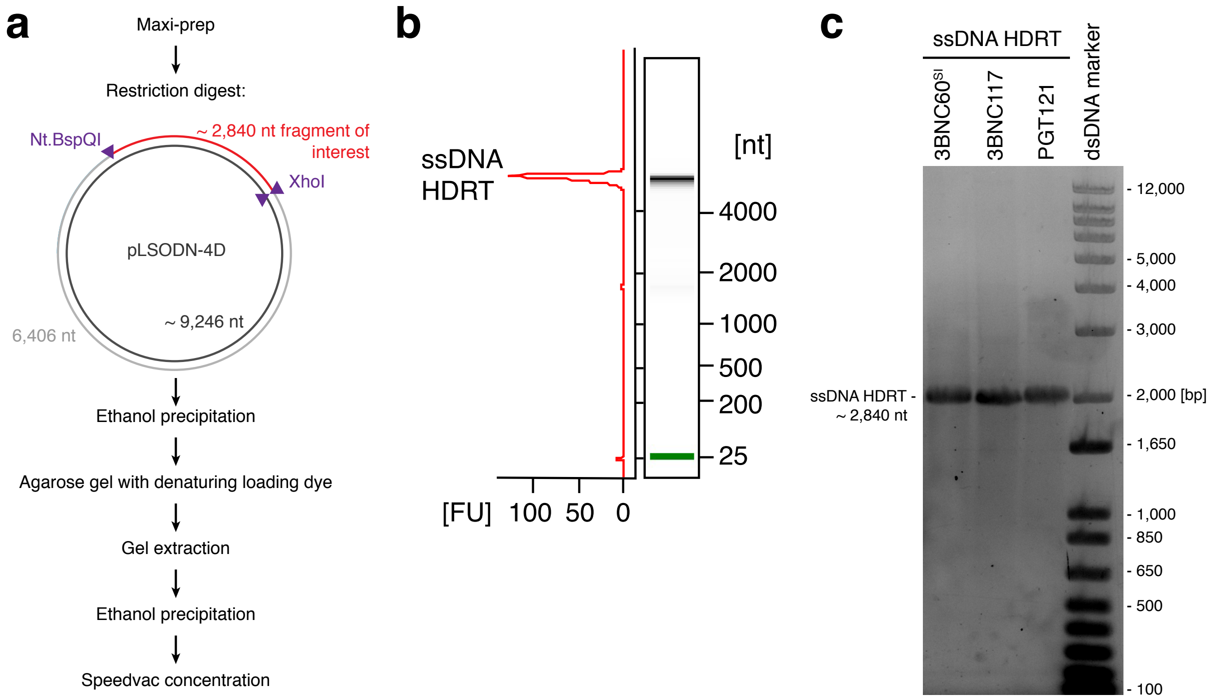


**Supplementary Figure 3: ssDNA HDRT template production**

**(a)** Flow chart of ssDNA production. HDRT templates were cloned into pLSODN-4D, Maxi-prepped, sequence verified and digested with restriction enzyme *XhoI* and the nicking endonuclease *Nt.BspQI* to produce 3 ssDNA fragments of the vector. Denaturing loading buffer was used to separate the 3 fragments by conventional agarose gel electrophoresis as indicated. ssDNA HDRT quality and integrity was verified using **(b)** Bioanalyzer and **(c)** agarose gel electrophoresis.

Supplementary Fig. 4

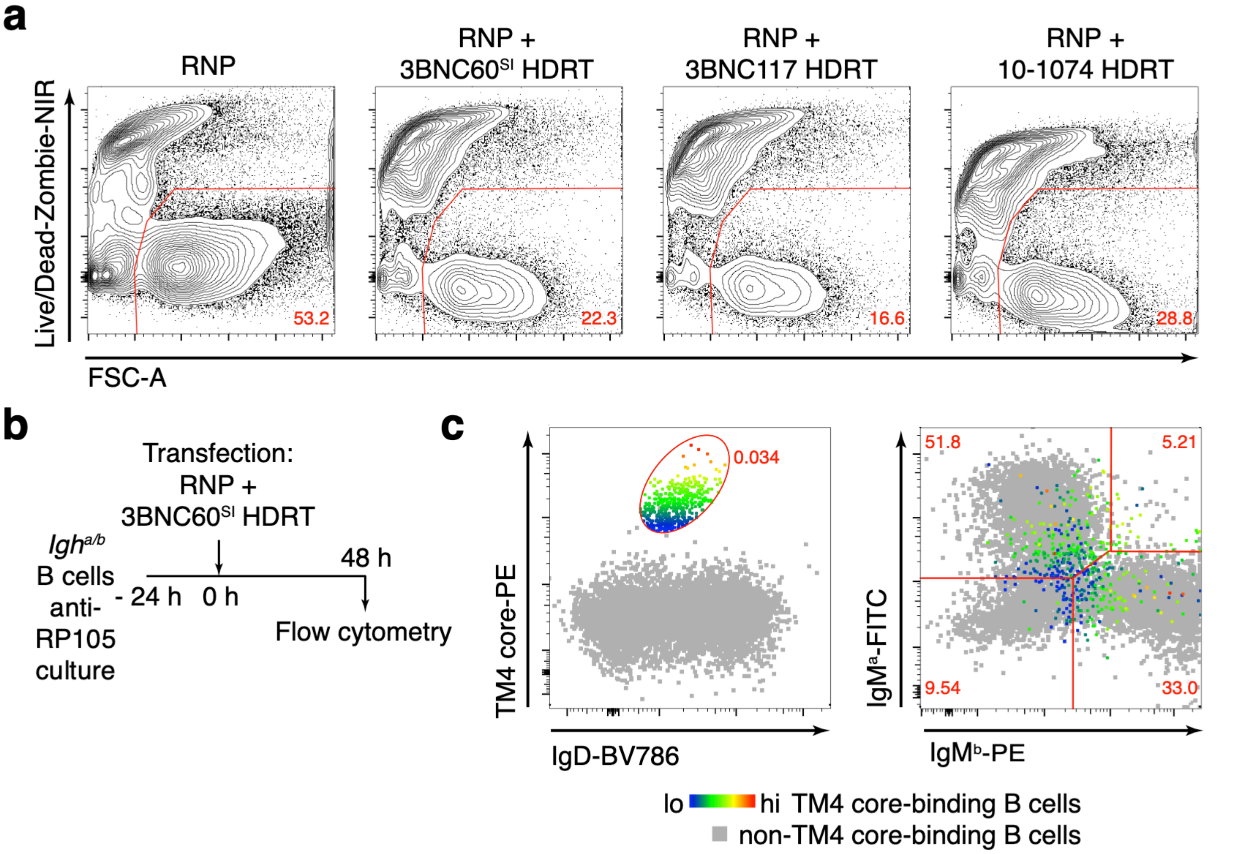


**Supplementary Figure 4: Cell viability and allelic exclusion of bNAb expressing murine B cells.**

**(a)** Flow cytometric plots showing percentage of live cells among all events 48 h after RNP ± HDRT transfection. Related to Fig.2b,c**. (b)** Experimental set up for (c). **(c)** Overlays of flow cytometric plots of TM4 core binding cells and non-binding B cells, both pre-gated on $\lambda^-$ B cells. TM4 core mean fluorescence intensity (5.89 x $10^3$ to 1.28 x $10^5$) is color mapped onto TM4 core-binding cell population. Numbers represent the percentage of TM4 core-binding cells among $\lambda^-$ B cells (left) or the percentage of TM4 core-binding B cells in the respective gate (right). Concatenate of 5 technical repeats in 2 independent experiments are shown.

Supplementary Fig. 5

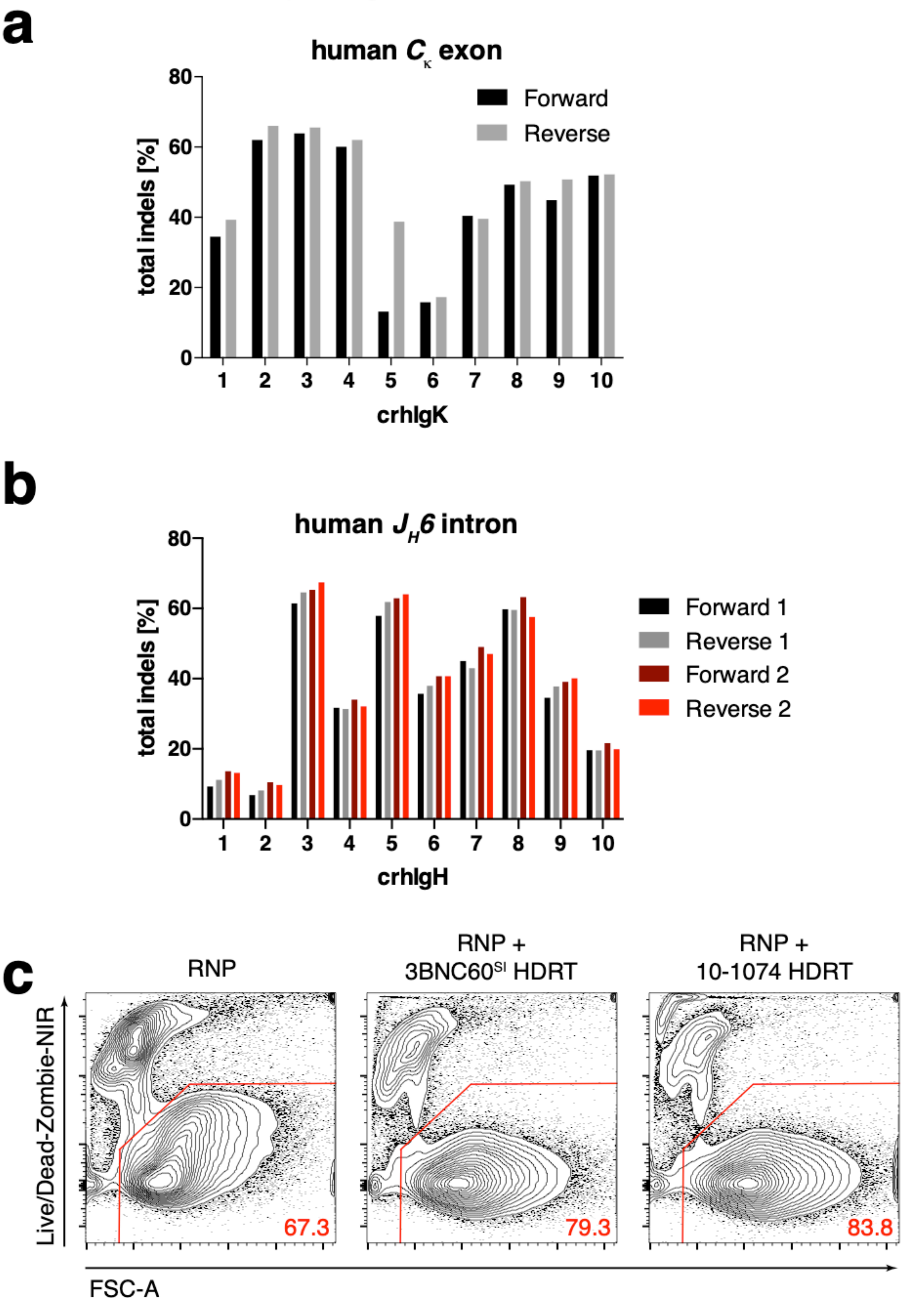


675

**Supplementary Figure 5: TIDE analysis and viability of primary, human B cells after transfection.**

**(a)** TIDE assay 42 h after transfection, comparing the efficiency of creating indels of crRNAs targeting the human *IGKC* exon and **(b)** TIDE assay using 2 different primer sets, 24 h after transfection, comparing the efficiency of creating indels of crRNAs targeting the human *IGHJ6* intron. Forward/reverse indicate sequencing with forward/reverse primers respectively. Representative of 1 - 2 independent experiments per crRNA. **(c)** Flow cytometric plots showing percentage of live cells among all events 72 h after RNP ± HDRT transfection. Related to Fig.4d. Representative plots of 2 independent experiments are shown.

### Supplementary Fig. 6

**a**

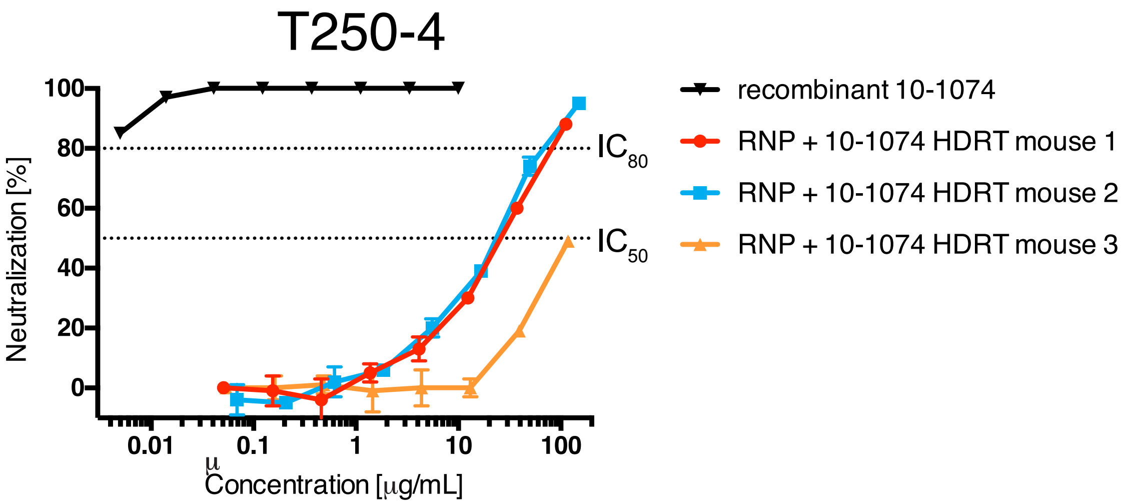


**b**

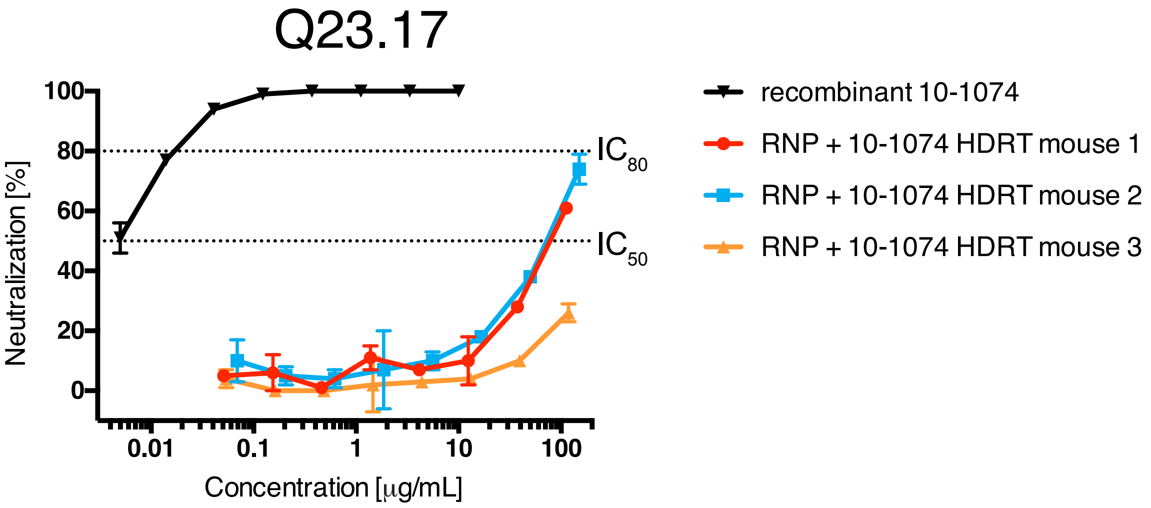


**c**

| | Titer in TZM.bl cells (μg/mL) | | | |
|---|---|---|---|---|
| **Virus** | **T247-23** | | **62357.14.D3.3489** | |
| **Sample** | **$IC_{50}$** | **$IC_{80}$** | **$IC_{50}$** | **$IC_{80}$** |
| RNP + $3BNC60^{SI}$ HDRT mouse 1 | >150 | >150 | >75 | >75 |
| RNP + $3BNC60^{SI}$ HDRT mouse 2 | >122 | >122 | >61 | >61 |
| RNP + $3BNC60^{SI}$ HDRT mouse 3 | >61 | >61 | >61 | >61 |
| recombinant 3BNC60SI (μg/mL) | **0.390** | **1.393** | **0.406** | **1.946** |

**Supplementary Figure 6: Serum neutralization of wild type mice adoptively transferred with edited B cells. Related to figure 4.**

**(a, b)** Neutralization curves for HIV strains T240-4 (a) and Q23.17 (b) of data summarized in Fig.4e of mice receiving 10-1074-edited B cells and immunized with cognate antigen 10mut. **(c)** HIV neutralization data of mice receiving 3BNC60$^{SI}$-edited B cells and immunized with cognate antigen TM4 core. Combined data from 2 independent experiments.
